# RingNet: An Interactive Platform for Multi-Modal Data Visualization in Networks

**DOI:** 10.64898/2026.01.20.700593

**Authors:** Liang Zhang, Xin Lai

**Author notes:** Corresponding author: Xin Lai, Faculty of Medicine and Health Technology, Tampere University, Tampere, Finland. Preprint: Liang Zhang, Xin Lai. RingNet: an interactive platform for multi-modal data visualization in networks. (2026). https://doi.org/10.64898/2026.01.20.700593.

## Abstract

The exponential growth of data in biomedicine has created an urgent need for intuitive visualization tools. These tools should effectively represent complex biological networks and remain accessible to domain experts without extensive computational training. Currently, there is a lack of a highly accessible, visually compact tool that can be used to examine multiple modalities simultaneously within arbitrary biological networks. Hence, we develop RingNet, a web-based interactive visualization tool that integrates computational efficiency with flexible, user-driven exploration. This tool addresses the community’s need to visualize multi-modal datasets within a single, compact network representation, as well as identify complex data patterns in network nodes. RingNet combines an R backend for network computation and layout optimization with a JavaScript-based viewer. Network topology and metadata are stored in JSON, while sample-level data are stored in indexed binary files for efficient retrieval and interactive browser-based visualization. It offers dynamic layout adjustments, node and edge filtering, and customizable color schemes for representing data. It can export reproducible, publication-ready figures in SVG and PNG formats. In our case studies, we use RingNet to visualize breast cancer patients’ omics profiles in a gene regulatory network and a cell-to-cell communication network in atopic dermatitis. This demonstrates RingNet’s ability to reveal biological relationships across multiple data modalities.

**Availability and Implementation:** The RingNet is available at https://ringnet.rd.tuni.fi and the source code and tutorial for RingNet is accessible at https://github.com/laixn/ringnet.

**Highlights:**

- RingNet enables intuitive, interactive visualization of multi-modal networks without requiring advanced computational or programming expertise.
- RingNet’s cores feature is its combination of flexible network input, compact multimodal node representations, and lightweight, web-based interactive visualization.
- RingNet has two functions: one to identify patterns in complex data at the node level, and the other to identify samples with homogeneous and heterogeneous profiles within network nodes.

Graphic abstract RingNet is a tool for visualizing multi-modal data in networks.

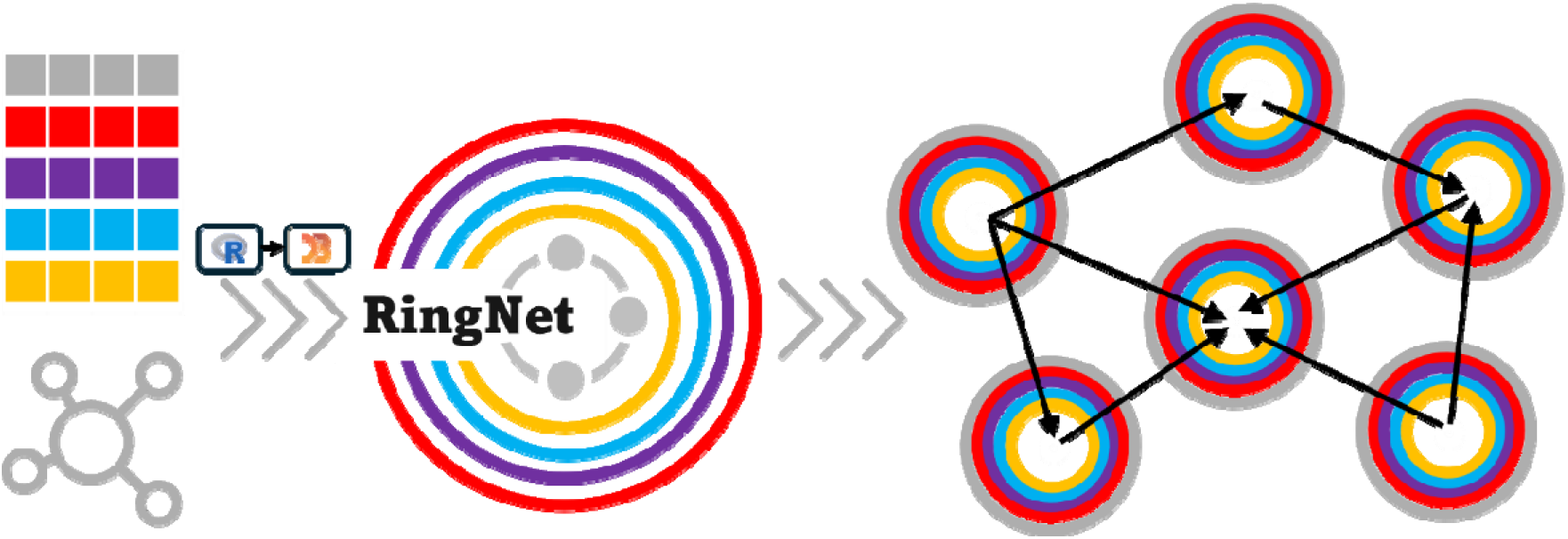

## Introduction

Network visualization provides an intuitive abstraction for complex data by encoding relationships between entities as edges connecting nodes. By emphasizing connectivity, dependency and topology, networks allow complex systems to be studied as structured, interacting units rather than isolated variables. In computational biology, network-based representations have therefore become central to modeling molecular interactions, regulatory dependencies and functional associations, supporting both exploratory visualization and integrative analysis (Barabási, Gulbahce, and Loscalzo 2011, Zitnik *et al*. 2024). Human biology operates across multiple hierarchical scales. Molecular components, including genes, transcripts, proteins, and metabolites, form pathways and regulatory networks that interact at the cellular, tissue, and organismal levels. High-throughput technologies now generate multi-omics datasets (e.g., genomics, epigenomics, transcriptomics, proteomics, and metabolomics) that capture this layered organization (Pinu *et al*. 2019). Interpreting such data requires analytical and visualization approaches that can preserve biological hierarchy while enabling navigation across molecular layers and contextual relationships (Schaffer and Ideker 2021, Kumar, Romano, and Ritchie 2025).

Encoding biomedical data in network-based representations enables the incorporation of these relationships into a wide range of big data tasks, including data integration (Valous *et al*. 2024, Baião *et al*. 2025), module detection (Lazareva *et al*. 2021, Gézsi and Antal 2024), and mechanistic interpretation (Schulte-Sasse *et al*. 2021, Lai *et al*. 2022) that are naturally fused with graph machine learning methods. Network visualization tools allow quantitative omics measurements to be overlaid onto interaction graphs, facilitating the identification of coordinated subnetworks and key regulatory elements. General-purpose platforms such as Cytoscape serves as foundational infrastructure for biological network visualization, offering an extensible environment for integrating diverse omics data with molecular interaction networks (Shannon *et al*. 2003). Cytoscape has a big community that develops different applications for enhancing network visualization. For example, Omics Visualizer allows network nodes to display different data sets, but only one value per set (Legeay *et al*. 2020). enhancedGraphics can convert network nodes into various types of charts, such as pie and bar charts, further enhancing Cytoscape’s visualization capabilities.

In addition to the Cytoscape ecosystem, there are other tools that serve a similar purpose for network visualization. VANTED is the earliest java-based tool for network visualization in systems biology (Junker, Klukas, and Schreiber 2006) and its upgrade includes a manifold support of systems biology standards like SBGN and SBML for visualization and data exchange (Rohn *et al*. 2012). Web-based platforms such as OmicsNet emphasize interactive and scalable visualization, including three-dimensional network layouts that facilitate the integration of heterogeneous molecular interactions and multi-omics data (Zhou *et al*. 2022). PaintOmics is also a web-based tool for visualizing multi-omics data using predefined pathways in biological databases like KEGG and Recatome (Liu T *et al*. 2022). In addition, several tools are developed to address specific challenges in multi-omics network visualization. NaviCom focuses on pathway-centric visualization by projecting multi-omics measurements onto curated signaling and metabolic maps, enabling intuitive “network portraits” of molecular perturbations (Dorel *et al*. 2017). MiBiOmics targets interactive exploration and multilayer network inference through a user-friendly Shiny-based interface, lowering the barrier for integrative multi-omics analysis but cannot provide network visualization (Zoppi *et al*. 2021). More recently, Arena3D supports hierarchical visualization of graphs using super nodes to connect different networks (Kokoli *et al*. 2023). SmCCNet 2.0 extends its network inference function by integrating phenotype information with multi-omics network construction and interactive visualization, facilitating the discovery of disease-associated subnetworks (Liu W *et al*. 2024). Complementary efforts focus on enhancing the visualization of complex, heterogeneous outputs rather than network inference itself. Transomics2cytoscape enables layered, trans-omics network visualization within Cytoscape, explicitly representing relationships between molecular layers in stacked network layouts (Nishida *et al*. 2024). Visualization-centric frameworks, such as MINGLE (Coletti *et al*. 2025) and aplot (Xu *et al*. 2025), emphasize coordinated, compositional views that integrate network-derived summaries with heatmaps, genomic tracks or other plot types, supporting comparative and contextual interpretation of multi-omics data.

Despite substantial progress, existing tools usually optimize specific data modalities, visualization metaphors, or analytical workflows. As an alternative, they visualize cross-omics as separate networks and then combine them using prior knowledge, such as information about molecular interactions. Therefore, a tool capable of visualizing diverse multi-omics datasets within a single, compact network representation is needed. To address this gap, we develop the web-based tool RingNet. Compared to available network visualization tools, RingNet’s defining feature is its combination of flexible network input, compact multimodal node representations and lightweight, web-based interactive visualization (Table 1). It offers the community a highly accessible and visually compact way of examining multiple modalities simultaneously on an arbitrary biological network.

**Table 1.**
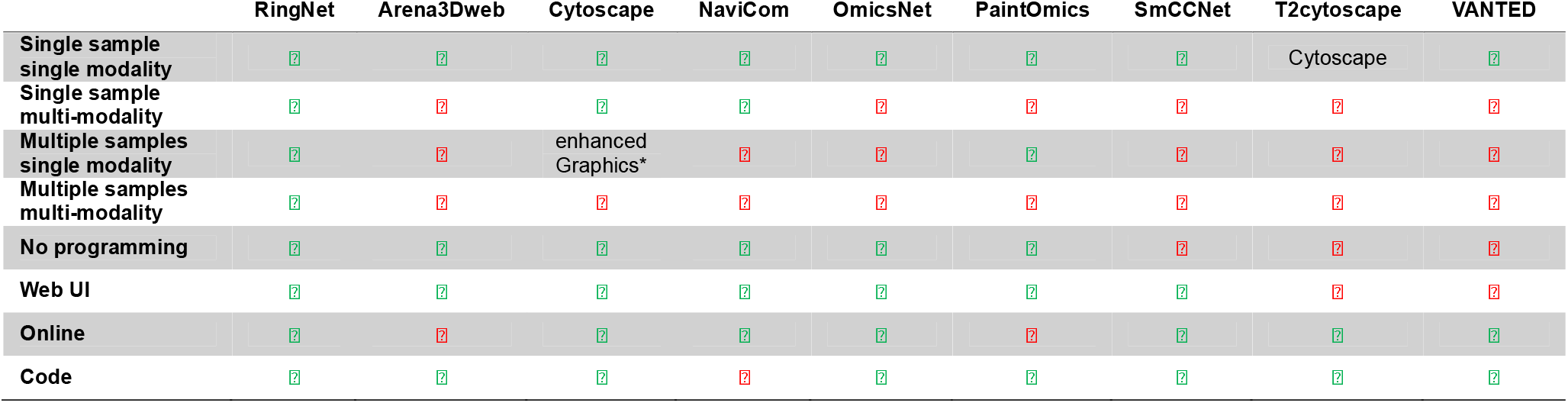
Comparison of RingNet’s functions with other tools. Symbols used for feature evaluations with ‘√’ for present, ‘√’ for absent. Texts indicate the applications or tools that are needed for a feature. The first four rows contain RingNet’s main visualisation features, while the rest comprise other basic functions. Single sample with single modality: A network node displays one data type for one sample. Each ring shows the data for one sample only. Single sample with multi-modality: A network node shows multiple data types for one sample. Each ring shows one data type. Multiple samples with single modality: A network node shows a single data type for many samples. Each ring shows one data modality, and each section of a ring shows data from a single sample. Multiple samples within multi-modality: A network node shows multiple data types for many samples. Each ring shows one data modality, and each section of a ring shows a single sample’s data. No programming required: Whether the user needs programming skills to use the tool. Web UI: Whether the tool has a web-based user interface (UI). Online: Whether the tool’s website is still working. Code: Whether the tool’s code is open access. VANTED: https://cls.uni-konstanz.de/software/vanted. Cytoscape: https://apps.cytoscape.org. OmicsNet: https://www.omicsnet.ca/OmicsNet/home.xhtml. PaintOmics: https://paintomics.uv.es. NaviCom: https://navicom.curie.fr/bridge.php. SmCCNet: https://smccnet.shinyapps.io/smccnetnetwork. T2cytoscape (Transomics2cytoscape): https://github.com/ecell/transomics2cytoscape. Arena3Dweb: http://bib.fleming.gr/Arena3D. *: enhancedGraphics is not running in the latest version of Cytoscape anymore. A more detailed and elaborative comparison can be found in SM Excel Table S1.

|  | RingNet | Arena3Dweb | Cytoscape | NaviCom | OmicsNet | PaintOmics | SmCCNet | T2cytoscape | VANTED |
|---|---|---|---|---|---|---|---|---|---|
| **Single sample<br>single modality** | ✓ | ✓ | ✓ | ✓ | ✓ | ✓ | ✓ | Cytoscape | ✓ |
| **Single sample<br>multi-modality** | ✓ | ✗ | ✓ | ✓ | ✗ | ✗ | ✗ | ✗ | ✗ |
| **Multiple samples<br>single modality** | ✓ | ✗ | enhanced<br>Graphics* | ✗ | ✗ | ✓ | ✗ | ✗ | ✗ |
| **Multiple samples<br>multi-modality** | ✓ | ✗ | ✗ | ✗ | ✗ | ✗ | ✗ | ✗ | ✗ |
| **No programming** | ✓ | ✓ | ✓ | ✓ | ✓ | ✓ | ✗ | ✗ | ✗ |
| **Web UI** | ✓ | ✓ | ✓ | ✓ | ✓ | ✓ | ✓ | ✗ | ✗ |
| **Online** | ✓ | ✗ | ✓ | ✓ | ✓ | ✗ | ✓ | ✓ | ✓ |
| **Code** | ✓ | ✓ | ✓ | ✗ | ✓ | ✓ | ✓ | ✓ | ✓ |

Technically, RingNet combines an R-based backend that transforms networks and multi-modal data into JSON-formatted graph objects and a JavaScript-powered web frontend that uses the JSON file to create interactive visualizations that users can customize. The tool’s functions include pattern identification, network decomposition for scalability, sample-based alignment for different types of data, and an interchange format that enables direct, attribute-driven visual mapping. Our goal in developing RingNet is to bridge molecular layers, preserve biological context, and support scalable, interactive exploration. This complements the visualization tools currently available for network biology and network medicine research.

## Methods

### RingNet’s design and architecture

RingNet is a three-tier architecture (Supplementary Figure S1). **(I) The layer for managing users’ sessions.** Users can upload a ZIP archive or multiple CSV or TSV files. The server then validates the inputs, creates a unique session identifier, stores the uploaded assets in a session directory, and sends back a JSON payload that describes the file paths and session metadata. **(II) The backend for data processing and export.** The core function of RingNet reads the interaction graph (edges and nodes), group membership, and one or more data types, such as omics data. The program harmonizes samples and features across inputs, constructs group-induced subgraphs, computes network statistics and two-dimensional layouts using R, attaches data vectors and standardized variants, and exports a JSON file and a binary file. The R backend stores topology, identifiers, and node-group metadata as indexed JSON. It also writes sample-level numeric measurements to a separate binary file. Each node and numeric layer stores its raw measurements as little-endian, 64-bit floating-point values (Float64), along with five statistics necessary for reconstructing the display scales. **(III) The frontend for interactive visualization.** The frontend uses JavaScript to load the JSON file and read and decode the binary file from the back end. It then renders group networks with interactive controls. The values of multi-modal data are mapped to visual channels through node and edge attributes, including size, color, directionality, curvature, and ring encodings. The program has export functions and supports SVG, PNG, and JSON output.

The PHP API invokes the R backend for the current session and reports the run status to the viewer. Once processing is complete, *upload_index.php* provides the session index, as well as the selected group’s topology and metadata, all in JSON format. For each request for a batch of nodes, *upload_index.php* uses the group’s starting offset in the JSON index, the requested node range, and the fixed size of each node record to calculate the byte offset and length in the binary file. Then, it streams the requested bytes to the browser. *ringnet_binary.js* decodes these bytes into numeric values and attaches them to the corresponding nodes.

### Frontend configuration

The RingNet frontend viewer is a browser-based visualization client implemented with Sigma JS. Its primary responsibility is to consume the RingNet JSON artifact and binary file produced by the backend, support interactive exploration, visual encoding configuration, and export SVG and PNG files (Supplementary Figure S2). The viewer is designed to minimize the browser workload, so intensive processing (such as subgraph extraction, data normalization, and JSON artifact preparation) is handled by the backend while users focus on rendering and interaction. Specifically, the viewer first loads the JSON index for the current session and retrieves the topology and metadata for the selected network group. Next, it calls ringnet_binary.js to request sample-level data in batches of consecutive nodes. For each request, *upload_index.php* looks up the group’s starting offset in the JSON index. It uses the requested node range and the fixed size of each node record to calculate the byte offset and length in the binary file. The PHP endpoint then streams the requested bytes to the browser. Finally, *ringnet_binary.js* decodes the bytes into numerical values and attaches them to the corresponding nodes in JSON node order.

The viewer uses Sigma.js and WebGL2 to render the network, with a 2D canvas overlay for the node rings. Users can filter nodes and edges, adjust color mappings, and zoom. They can also switch between raw and normalized values, such as min-max normalization and z-scores. Users can export the current viewport as a PNG or SVG file, or save the network and view settings as a JSON session snapshot for reuse.

### Deployment and security configuration

RingNet uses Apache as a web server and reverse proxy to securely and uniformly provide access to the application (Supplementary Figure S3). Apache handles Transport Layer Security (TLS) termination, Hypertext Transfer Protocol Secure (HTTPS) enforcement, security headers, and access control. It also routes both static viewer assets and API requests under a single domain. Backend services remain bound to internal ports, which improves security isolation. JSON artifacts are served through authorized API endpoints to prevent unauthorized access and directory traversal. HTTPS is enforced via port 443, with redirects from port 80, as well as with optional HTTP Strict Transport Security (HSTS). This architecture simplifies user configuration, improves performance through user-side rendering, and supports reproducibility via session-scoped artifacts and exportable session state.

### The implementation of network visualization in RingNet

RingNet requires two mandatory input tables provided in comma-separated values (CSV) format (Figure 1). (**I**) **Graph edges**. The edge table provides interactions within networks. It must contain two columns that specify the source and target nodes of each interaction. An optional weight column is supported to include qualitative or quantitative information, such as the correlation coefficient of the interacting nodes. If the weight column is absent, the weights are initialized to the default value of 1. Optional annotation fields, such as interaction type or interaction identifier, are preserved and propagated to the output when present. (**II**) **Graph nodes**. The node table defines network nodes and must contain a unique identifier or symbol for each one.

**Figure 1.**
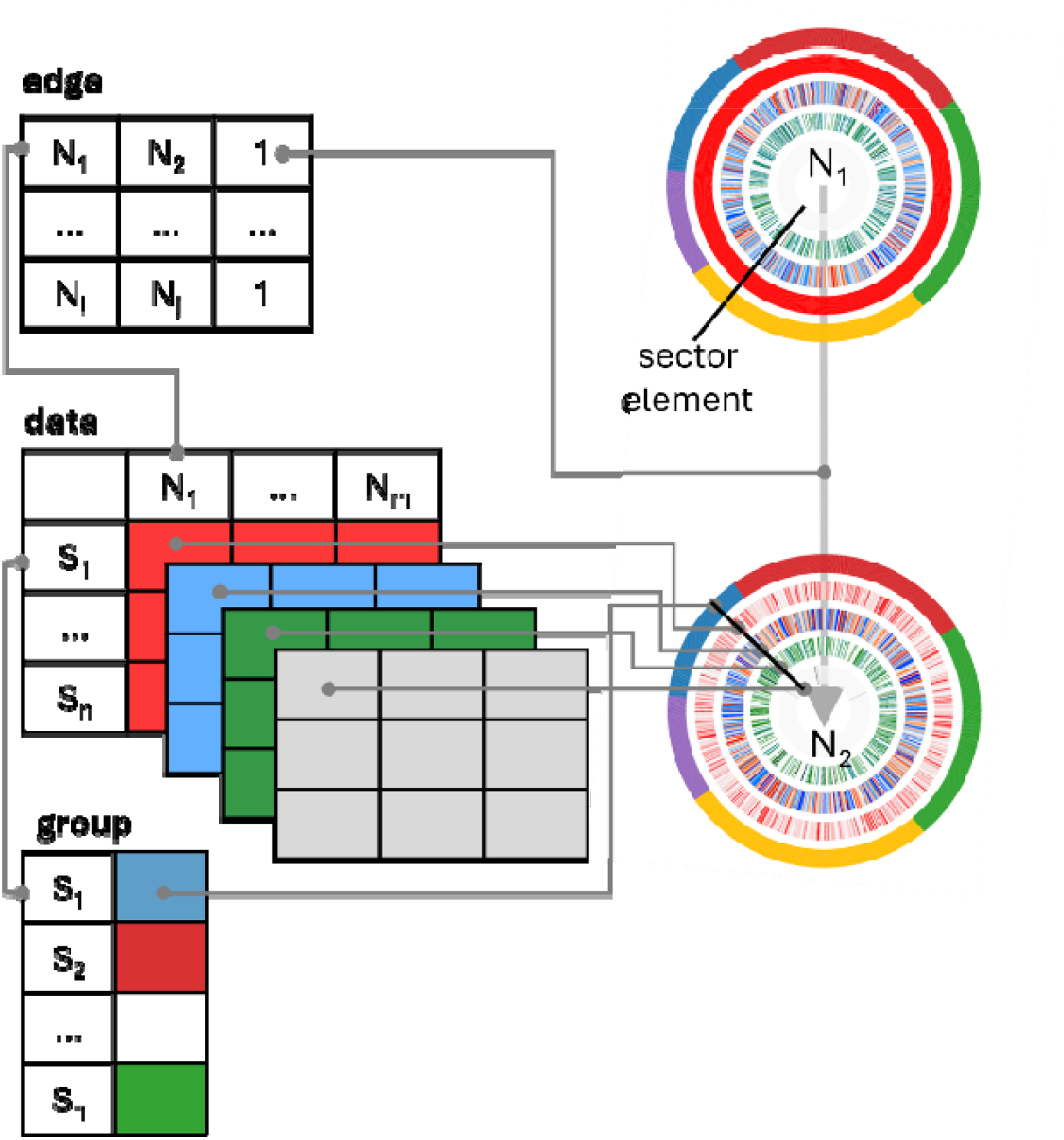
RingNet’s visualization protocol. RingNet can integrate up to four numerical data sets and one categorical data set into a node. Each data matrix has a structure in which the rows represent sector elements (e.g., samples) and the columns represent network nodes (e.g., genes). In a node, each ring represents one data matrix, with the categorical matrix on the outside and the numeric matrices inside. Each ring contains the same number of sector elements and data from the same sample are shown on the aligned segment sectors on all five rings, enabling the joint visualization of multi-modal data within a single node. The edge table shows the interactions between nodes (e.g., gene-gene interactions), which can be assigned arbitrary (e.g., 0 or 1) or numerical weights (e.g., correlation coefficients). The gray lines indicate the sources of the data that RingNet uses to visualize networks and the shared identifiers of different data that enable cross-modal alignment.

The group table file (**Node group**) assigns each node to a single group. The table must contain a group column and an identifier column for network nodes. Duplicate identifiers are removed to enforce a one-to-one mapping between nodes and group assignments. This file allows users to simultaneously visualize multiple networks. Additionally, RingNet supports up to five data matrices with continuous, integer, or categorical values in CSV format. Each matrix should have rows that correspond to the sector elements of a ring diagram and columns that correspond to the nodes of a network. Row names are treated as identifiers for sector elements and are used for cross-modal alignment. RingNet harmonizes heterogeneous inputs through intersection-based alignment rules defined at the level of sample and node identifiers. These rules guarantee consistency of indices across multiple modalities without imposing assumptions of numerical comparability between heterogeneous data types.

### Network construction and layout at the backend

The network is constructed using the R package *igraph*. By default, the network is regarded as directed, but it can also be set as undirected. When treating the network as directed, node connectivity summaries may include total degree, in-degree, and out-degree. For undirected graphs, only total degree is used. RingNet computes two-dimensional node coordinates for each network using stress-based layout. The algorithm treats edges as elastic constraints between nodes. Nodes that are directly or strongly connected are placed closer together. In contrast, nodes that are connected only indirectly or separated by intermediate nodes are positioned further apart. Consequently, the layout expands the overall graph structure so that densely connected nodes become compact and sparsely connected or weakly interacting nodes are pushed further apart. The coordinates are scaled by a constant factor to improve visual separation. If the stress-based layout optimization fails, as in the case of very small or fragmented networks, RingNet uses a ForceAtlas2 layout. The resulting coordinates are exported as fixed node attributes, which enables deterministic frontend rendering without requiring users to compute the layout.

### Network visualization at the frontend

RingNet exports visualization-ready JSON groups containing nodes and edges with precomputed layouts, identifiers, multi-modal data to support efficient interactive network rendering. In the frontend visualization, each node in the network can have up to five rings, each representing a data set. For numeric data, RingNet provides raw, min-max normalization, and z-normalization options to enable flexible frontend visualization. RingNet also provides raw, min-max normalized, and z-normalized weights for visualizing edges in the network. Node labels are identifiers (e.g., gene symbol, patient sample id, and cell type). Users can filter nodes and edges by their containing values and topology-derived properties such as node degree, and customize their aesthetic properties, such as node and edge colors. At the node level, RingNet introduces multi-modal concentric ring glyphs. Each node is represented by a set of concentric rings, with each ring representing a data modality. Samples are mapped to angular sector elements that are shared by all rings to ensure cross-modal alignment. Color scales encode continuous values, discrete palettes represent categorical data, and range-based selection allows for highlighting without removing data. Edges are rendered with flexible encodings that support multiple weight metrics, signed color mappings, and optional coloring by interaction type.

### Multiple network visualizations

RingNet also supports multiple network visualization. Specifically, RingNet uses a process-based cluster with a configurable number of CPU cores, exports the necessary objects to the workers, and applies over the network identifiers. This design leads to near-linear speedups when network sizes are balanced, and memory overhead is manageable. The main trade-off is the replication of data matrix and edge table memory across multiple CPUs. Under the current configuration, RingNet can simultaneously compute and visualize up to 25 moderately sized networks, (ranging from 52 to 1,872 nodes and from 20 to 6,100 edges) within several minutes (Supplementary Table S1). Users can select individual groups for interactive exploration in the viewer. Processing the largest network requires approximately 21 seconds and reaches a peak memory consumption of about 400 MB, with memory usage increasing steadily during execution.

### Node-based pattern discovery

It is usually challenging to detect biologically meaningful patterns in multi-omics networks through visual inspection alone, especially when the number of samples is large. Therefore, we need a function that can reveal patterns hidden in the data. For example, in a cancer cohort, a gene may exhibit high expression accompanied by low DNA methylation in one subset of patients, while another subset displays the opposite pattern of low expression and high methylation. Such anti-correlated relationships are of particular biological interest because aberrant DNA methylation is frequently associated with transcriptional dysregulation in cancer (Imran *et al*. 2025). However, when hundreds of samples are represented as ring sectors within network nodes, it becomes difficult to determine whether these patterns are consistently observed across the cohort, restricted to specific patient subgroups, or associated with disease progression. To address this limitation, RingNet incorporates a pattern discovery function that systematically quantifies multi-omics deviation patterns and prioritizes candidate nodes with patterns of interest (see Supplementary Materials for details). Rather than relying solely on visual inspection, the method ranks samples and nodes according to their multi-omics profiles, enabling the identification of representative molecular patterns and subgroup signatures. This approach complements the visual exploration workflow by transforming complex multi-omics observations into interpretable and searchable network features, while preserving the flexibility of user-defined visual encoding, filtering, and data export. Importantly, users can adjust the node filtering thresholds, which affect only the selection and display of candidate nodes. This ensures a stable visual context for comparative analysis, as the network layout, node positions, edge geometry, and color settings remain unchanged.

### Sample selection based on multi-omics profiles

RingNet features a visualization mode that is designed to identify representative yet heterogeneous or homogeneous samples within predefined groups, such as tumor stages. Rather than randomly selecting samples or displaying them all simultaneously, this method first quantifies the multi-omics deviation strength of each sample. This quantification uses weighted z-score profiles derived from gene expression, DNA methylation, copy number variation, and somatic mutation data. The scoring framework considers global network-level deviations and local node-level differences. It also incorporates robust, percentile-based statistics that capture strong tail signals from highly perturbed genes. When tumor stage annotations are provided, samples are stratified by stage prior to selection of a representative sample. Within each stage, the algorithm first selects the sample with the highest overall deviation or similarity score. Then, it applies a farthest-first or closest-first strategy in the multi-omics feature space to select additional samples that differ from or are similar to those initially selected. This approach preserves intra-stage heterogeneity or homogeneity while preventing the overrepresentation of a single molecular pattern within a group of patients. Finally, RingNet recalculates robust node-level variation ranges from the selected samples and automatically retains nodes that exceed predefined thresholds in key omics layers, such as gene expression and DNA methylation. This gives biologically heterogeneous or homogeneous patterns within the network (see Supplementary Materials for details).

## Results

RingNet is a powerful tool that integrates multi-modal data into networks, enhancing the data visualization demanded by the biomedical group (Figure 2). The group generates complex data to characterize and understand human diseases such as cancer. For example, the cancer genome atlas (TCGA) project is the first to generate large-scale, multi-omics profiles for patients across 33 cancer types that enables systematic cross-cancer analyses (Cancer Genome Atlas Network 2012). Additionally, an increasing number of single-cell projects generate RNA-seq data for millions of cells to characterize tumor microenvironments (Yuan *et al*. 2019, Han *et al*. 2023), immunology (Yin *et al*. 2026), and autoimmune diseases such as atopic dermatitis (He *et al*. 2020). These complex data require a sophisticated tool for network-based visualization and integrative exploration to reveal relationships across molecular layers.

**Figure 2.**
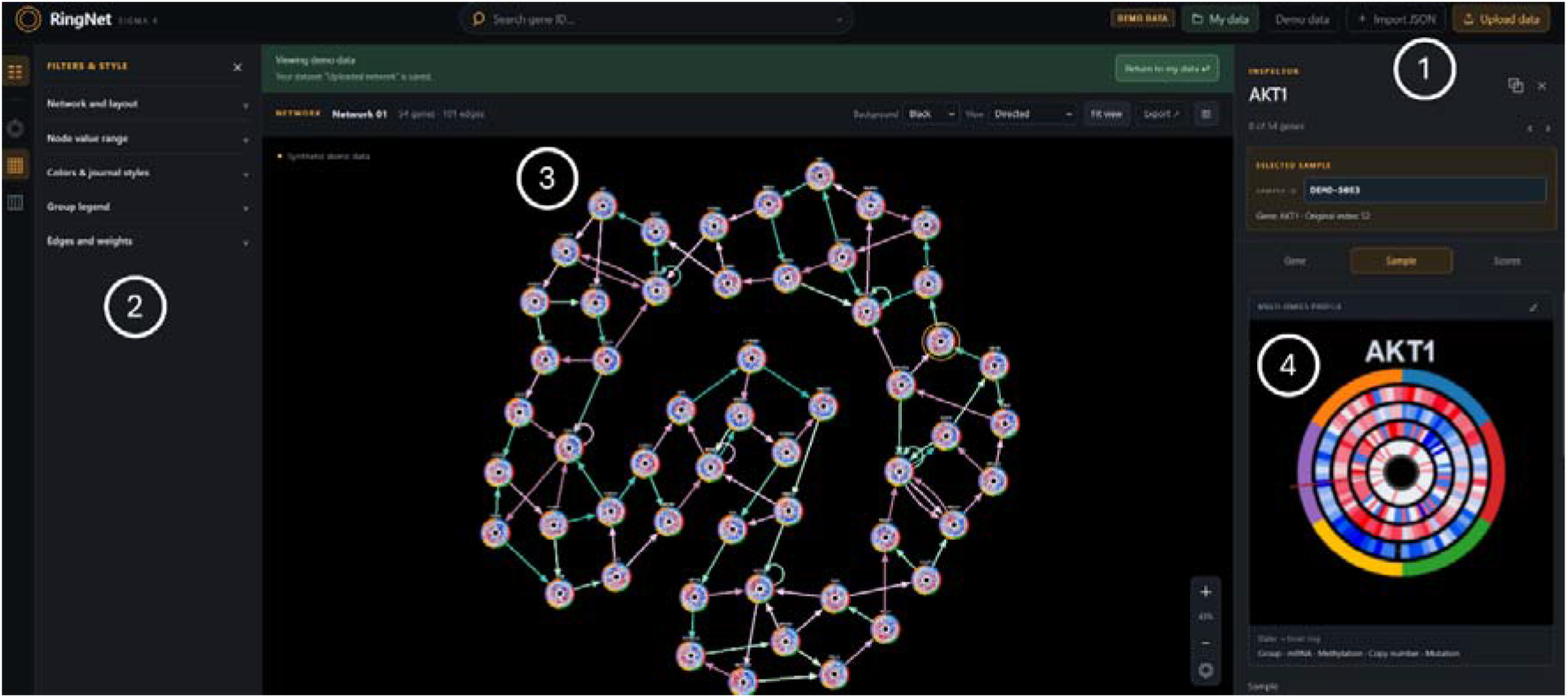
RingNet’s Viewer interface. (1) Users can access the tutorial, open the built-in demo dataset, upload CSV files via Upload data, or reopen a saved network via Import JSON. For CSV inputs, the R backend generates indexed JSON containing the network topology and metadata, along with a companion binary file containing sample-level values. The viewer combines these outputs to display the network. (2) The filters and style panel lets users adjust the layout, filter nodes and edges, and customize colors. Preset palettes are inspired by the visual styles of The New England Journal of Medicine, Nature Cancer, The Lancet, and Science. (3) The network canvas updates as users apply these settings and supports panning, zooming, and dragging nodes. (4) Selecting a node or a ring sector updates the inspector panel, which displays the selected node’s multi-data profile and provides access to sample details and scores. The detailed technical workflow of RingNet can be found in Supplementary Figure S4.

The primary function of RingNet is to integrate different data sets into a network node. These data sets can contain numerical or categorical values, allowing for the combination of different types of information. Aligned data for a node’s sector elements (representing a patient or cell) makes interpreting results more convenient. Matching the profile of one object allows information to be associated, thereby enriching the information hierarchy and ensuring better explanations and interpretations. RingNet’s comprehensive collection of functionalities establishes it as a robust tool for creating composite networks, empowering researchers to produce intuitive data visualizations. RingNet enables effective communication and in-depth interpretation of network science through precise control and annotation of elements. RingNet also provides a function for the parallel visualization of multiple networks, which facilitates the creation of a series of networks for comprehensive analysis, such as pathway networks from gene set enrichment analysis (Subramanian *et al*. 2005). Specifically, users can visualize multiple networks of identified enriched pathways in a single run. This function enables users to efficiently generate visualization results from large-scale data analyses. RingNet also provides two major functions for exploring networks enriched with multi-modal data (see Methods). The **sample-identification** function identifies representative heterogeneous or homogeneous samples using multi-modal data. It can select group-specific samples that maximize diversity or similarity in network nodes. The **node-based pattern discovery** function systematically detects multi-modal patterns in network nodes, such as anti-correlations between gene expression and DNA methylation in a large cancer patient cohort (Figure 3). By filtering candidate nodes with a particular pattern, it reveals subgroup-specific and progression-associated molecular signatures while preserving a stable network visualization context. Furthermore, since RingNet supports uploading JSON files for computed networks, users do not need to rerun the time-consuming tool to reproduce the results. Instead, they can use the JSON file for their network. RingNet can then visualize the results directly. These functions can save users significant time when visualizing massive networks or repeatedly refining the final network output. The web-based RingNet makes it a suitable tool for researchers with or without programming skills.

**Figure 3.**
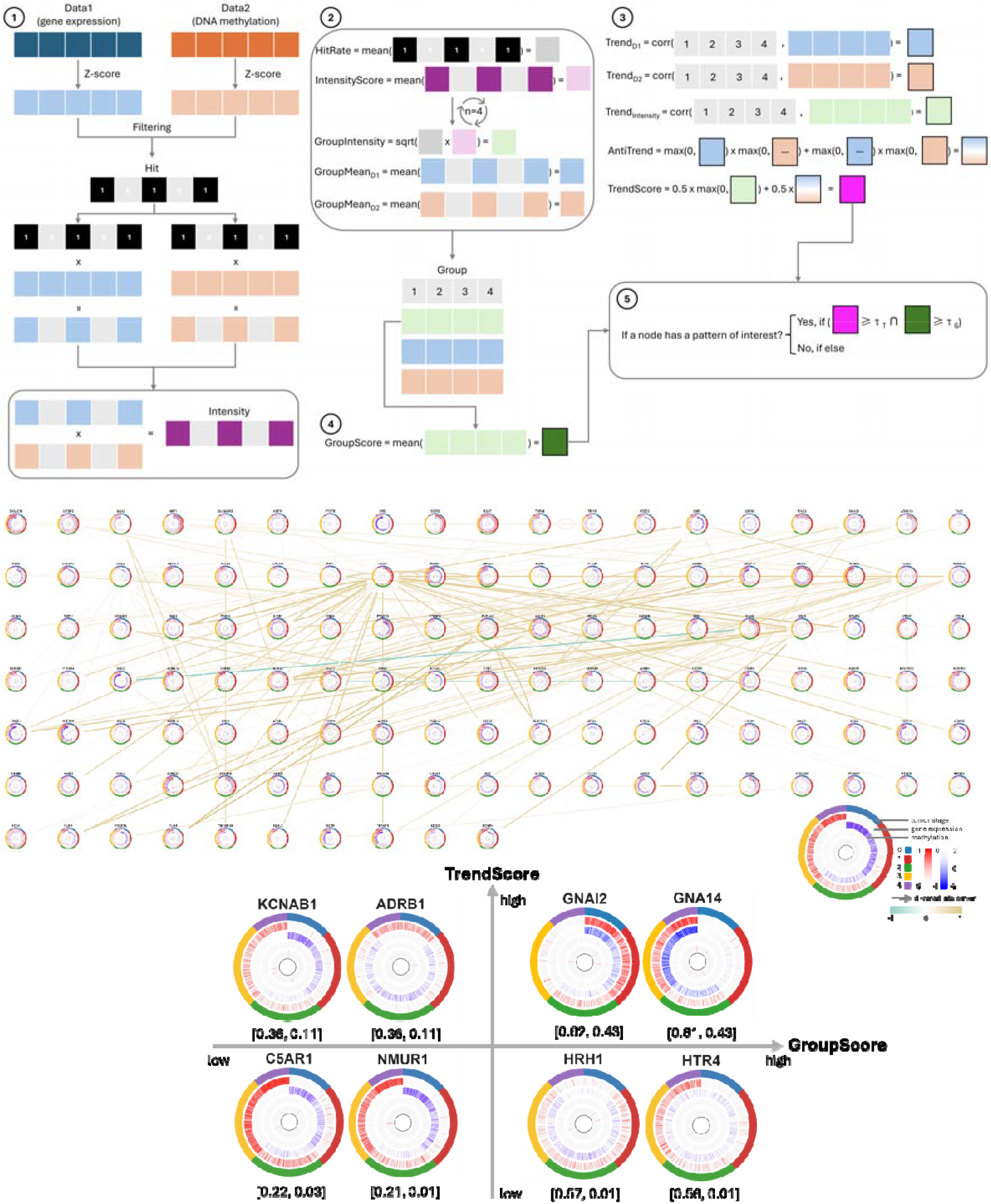
Illustration of the node-based pattern discovery function. (**Top**) The pattern discovery function operates through five sequential steps. First, sample-level pattern hits are identified based on user-defined high and low thresholds. Hit intensity is quantified using z-score transformation of data. Second, when group annotations are available, hits are aggregated into group-level intensity scores. Third, TrendScore measures whether pattern evidence strengthens across groups. Fourth, GroupScore quantifies group-level support. Fifth, a two-threshold rule jointly selects nodes fulfilling the predefined criteria for GroupScore and TrendScore. (**Bottom**) The pattern discovery function quantifies the expression and methylation patterns of genes in a network. Some exemplar genes are grouped into four categories based on their corresponding GroupScore and TrendScore values, shown in brackets. C5AR1 and NUMR1 have low scores, indicating sparse alignment and weak anti-correlation patterns in patients. They demonstrate invisible negative correlations in their profiles. HRH1 and HTR4 have high GroupScore and low TrendScore, indicating densely aligned profiles with weak anti-correlation patterns. These genes show weak negative correlations between the two profiles. KCNAB1 and ADRB1 have moderate GroupScore and high TrendScore, indicating moderately aligned profiles with anti-correlation patterns in some, but not all, groups. Overall, they show weak negative correlations. GNAI2 and GNA14 have high scores, showing densely aligned profiles with anti-correlation patterns and strong negative correlations between the two profiles. Each ring has three data sets including the group information that represents five tumor stage, gene expression profile, and DNA methylation profile. The edges are color coded using the correlation coefficients for gene-gene interactions. The genes’ descriptions can be found in Supplementary Table S2.

### Large-network evaluation

We evaluated RingNet on a network with 7,093 input nodes and 48254 edges. After upload, the elapsed time from starting the run to receiving confirmation of server-side completion was 4 min 55 s. The viewer reported a graph layout time of 2,708 ms.

### Visualizing multi-omics data for the TCGA breast cancer cohort

Integrating and visualizing patients’ profiles is essential for understanding the complex relationships between multi-omics data and for unravelling patterns hidden in a large cohort (Figure 4). Here, we demonstrate how RingNet can effectively combine and display TCGA data for breast cancer patients, including genomic mutations, copy number variations, DNA methylation, transcriptomics, and clinical annotations (Cancer Genome Atlas Network 2012). We begin with a gene regulatory network for breast cancer that contains 8,185 genes and 48,534 molecular interactions derived from Omnipath (Türei *et al*. 2021). Next, we identify a subnetwork of 16 genes related to the GPCR pathway that modulate proliferation, survival, angiogenesis, invasion, and immune evasion in breast cancer (Arang and Gutkind 2020). Then, we use RingNet to map the omics data of 651 patients from TCGA and genes’ expression correlation coefficients onto the network’s nodes and edges, respectively.

**Figure 4.**
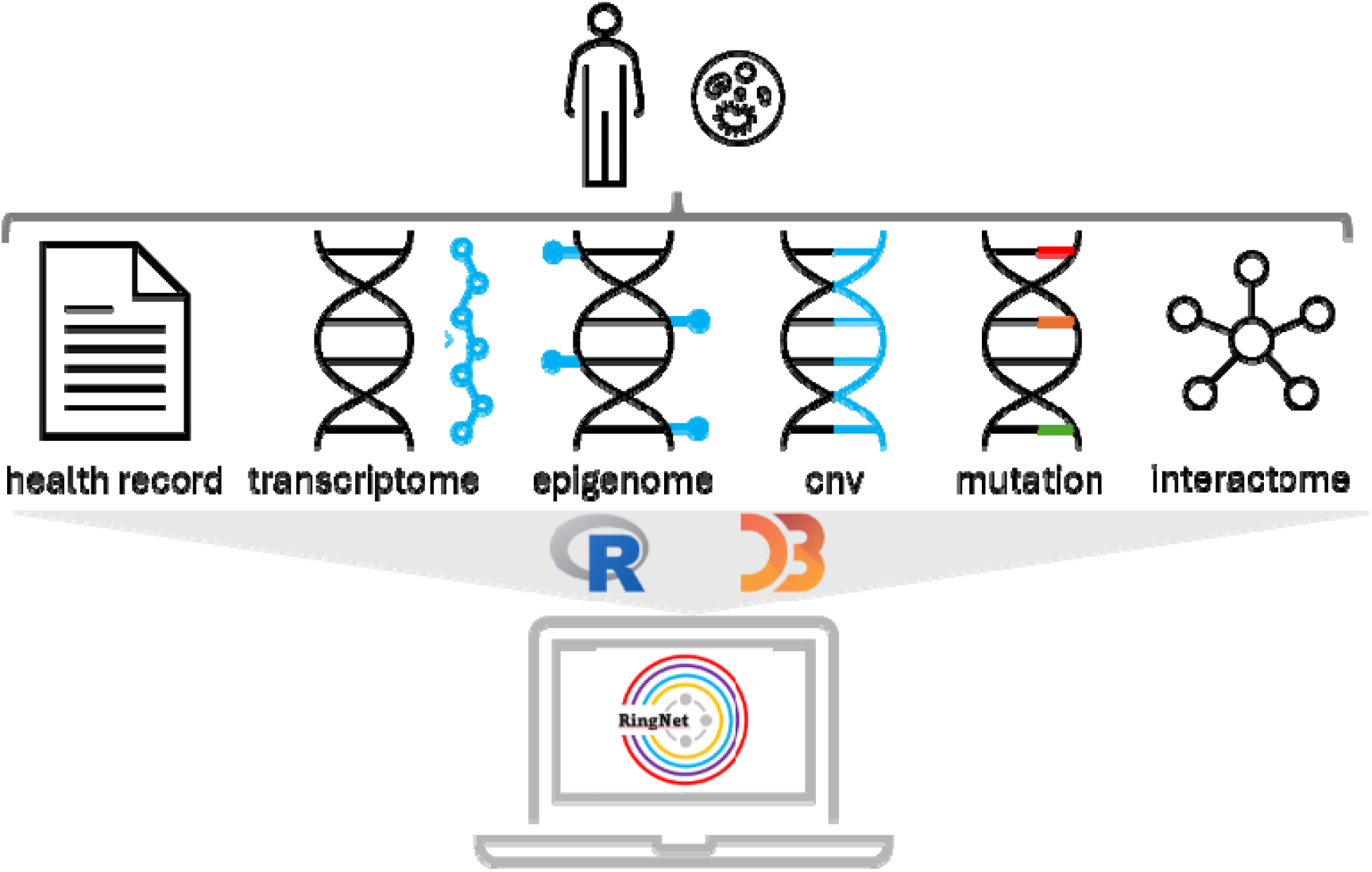
Multi-omics data integration in network visualization using RingNet. Multi-omics data, including DNA methylation, gene expression, somatic mutation, and copy number variation (CNV), as well as clinical records and molecular interactome data from a cohort study, are processed in R to build a network. This network is then exported as a JSON file that contains the omics profiles embedded within the nodes, as well as the network’s topology. The web interface uses HTML and D3.js to render the network, in which each node has a multi-ring layout displaying different types of data. Users can explore and refine the network through interactive operation.

As a result, we obtain a network comprising multiple layers of information (Figure 5). From the network, we can see whether patterns exist between the omics profiles of genes and the tumor stages of patients. It can also show whether genes that interact with a specific type of interaction, such as stimulation or inhibition, and quantify them using expression correlation coefficients. Some genes consistently exhibit high expression levels (e.g., GNAQ and GNAI2), some exhibit low expression levels (e.g., TACR1 and HRH2), and some exhibit mixed high and low expression levels (e.g., AGTR1 and GNB5). It can show correlation between specific molecular profiles, such as the correlation pattern between a gene’s expression and DNA methylation levels in the cohort. For instance, in the cohort, TACR1 typically exhibits low expression and methylation levels, whereas GNAI2 usually exhibits high expression and methylation levels. It can reveal whether patients have amplified or deleted copy number variations or mutations in certain genes. Most genes in the cohort exhibit copy number variation, and few gene mutations are observed. For example, most patients have amplified and deleted copy number variation in AGTR1 and GNAZ, respectively, but only two and one patient have mutations in these genes, respectively.

**Figure 5.**
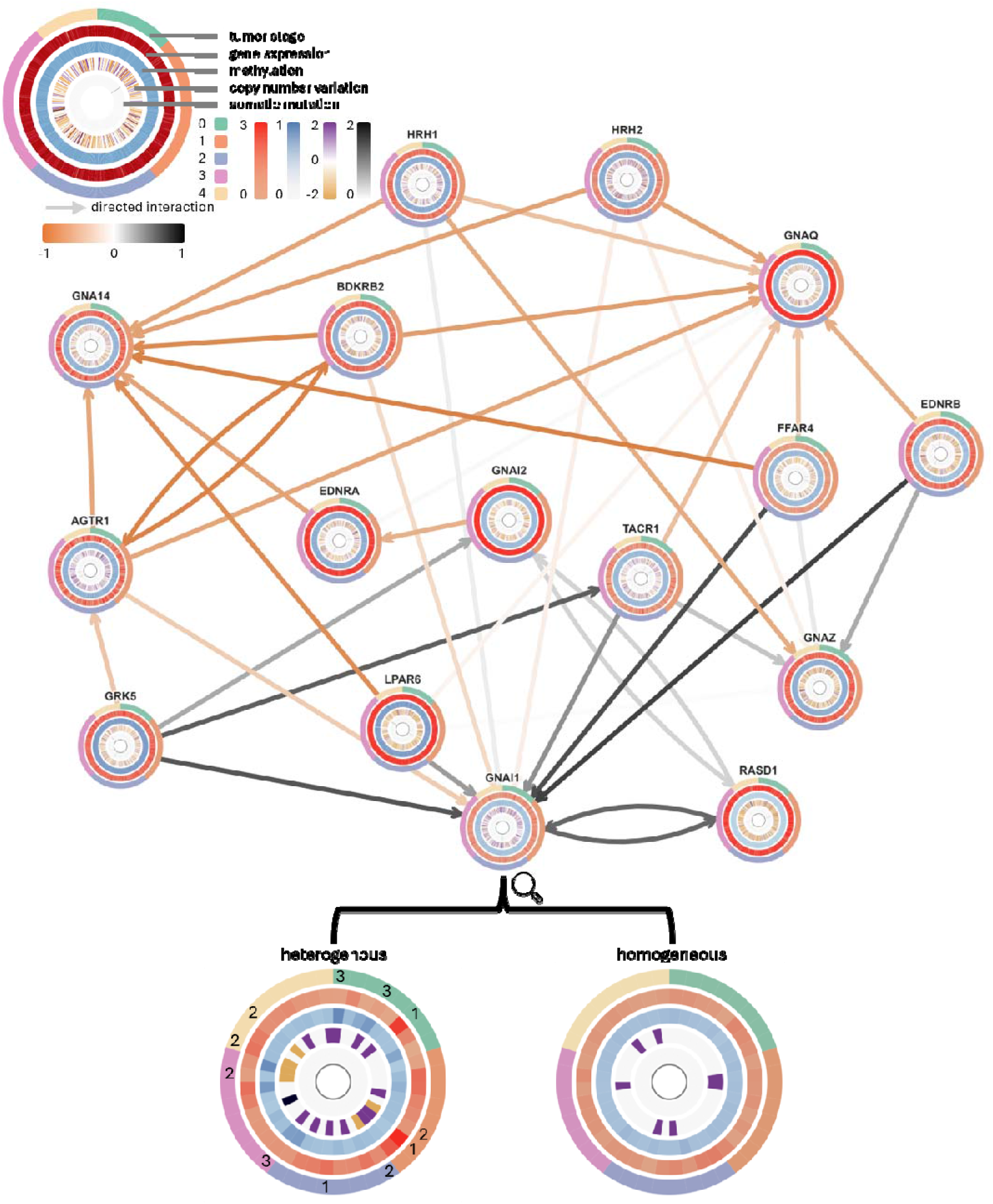
Gene regulatory network for breast cancer. Each node (i.e., gene) contains five rings that show omics data from the TCGA breast cancer cohort. From the outside in, the rings show the following: patients’ tumor stages, gene expression profile, DNA methylation profile, copy number variation, and somatic mutation profile. There are 468 sector elements in each ring, representing the corresponding profiles of 651 patients. The edges represent directed gene-gene interactions, and their colors indicate positive or negative expression correlation coefficients, respectively. The enlarged GNAI1 nodes illustrate genomic heterogeneity and homogeneity in different tumor stages. Genomic heterogeneity is indicated by numbers 1, 2, and 3, and genomic homogeneity is indicated by patients with consistent gene expression and methylation profiles in different tumor stages. A total of 32 patients are shown, eight for each tumor stage. The patients show the most distinguished profiles between each other in each tumor-stage. The genes’ descriptions can be found in Supplementary Table S2.

Multi-omics patterns can provide additional information when specific patient profiles are examined in more detail. For instance, stage-specific patterns can be observed across aligned patient groups. As tumor stages increase, GNAI1 shows stronger copy number variation signals. Gene expression and DNA methylation profiles appear relatively stable across stages. This suggests that genomic alterations contribute more prominently to stage-associated molecular heterogeneity in GNAI1 than transcriptional and epigenetic profiles do. Additionally, the aligned ring structure enables direct inspection of patients with heterogeneous or homogeneous molecular profiles within the same tumor stage (see Methods). For example, some patients (class 1) show strong GNAI1 expression, mild DNA methylation, and copy number amplification, while some patients (class 2) exhibit low GNAI1 expression, mild DNA methylation, and copy number deletion. This suggests that transcriptional activation is potentially associated with genomic gain. Some patients (class 3) display reduced GNAI1 expression accompanied by strong DNA methylation and copy number amplification, indicating that transcriptional inhibition is potentially associated with an epigenetic modulation independent of copy number status. For the identified patients with homogeneous molecular profiles across different tumor stages, they show consistent gene expression and DNA methylation profiles in GNAI1, and most of these patients have no copy number variations or mutations in GNAI1. These aligned multi-omics patterns demonstrate RingNet’s ability to reveal inter-patient heterogeneity and homogeneity, something that is impossible to achieve with the available independent tools or the tools embedded in Cytoscape. Taken together, RingNet allows for the effective integration of genomics data into a molecular network to uncover data pattern and variation in the cohort.

### Visualizing the cell-cell communication network for atopic dermatitis

Single-cell analysis emerges as a crucial tool in functional genomics with widespread applications in disease research. Key steps in single-cell transcriptomic analysis include cell clustering, identifying marker genes, and annotating cell types. These components are typically visualized collectively, often relying on existing tools that limit the integration of additional information. We demonstrate RingNet’s capabilities by applying it to scRNA-seq data from lesional human skin cells of patients with atopic dermatitis (He *et al*. 2020). This dataset contains information on 39,042 cells that comprise the microenvironment of this autoimmune disease. We use CellChat to infer the intercellular communication network between the identified cells. This tool models the probability of cell-to-cell communication by integrating gene expression with prior knowledge of interactions between signaling ligands, receptors, and cofactors, based on the law of mass action (Jin *et al*. 2021). Afterwards, we use RingNet to visualize the disease’s cell-cell communications (Figure 6). The network shows how different types of cells communicate in the microenvironment and the cells’ gene expression profiles. Specifically, inflammatory dendritic cells (Inflam. DC), conventional DCs (cDC1 and cDC2), T cells (TC, Inflam. TC, and CD40LG⁺ TC), Natural killer T cells (NKT), and Langerhans cells (LC) form dense communication hubs with fibroblast populations (Inflam. FIB, COL11A1⁺ FIB, FBN1⁺ FIB, and APOE⁺ FIB). Mechanistically, these interactions are mediated by ligand-receptor signaling, including cytokines, chemokines, and costimulatory molecules. Activated immune cells stimulate fibroblasts to produce extracellular matrix components, inflammatory mediators, and growth factors, while fibroblasts feedback to immune cells, sustaining immune cell recruitment, polarization, and tissue remodeling. This bidirectional signaling underlies epidermal thickening, fibrosis, and chronic inflammation typical of atopic dermatitis (He *et al*. 2020). Together, the results demonstrate the potential of RingNet to enhance visualization in single-cell data and advance our understanding of the underlying cell and molecular biology of atopic dermatitis.

**Figure 6.**
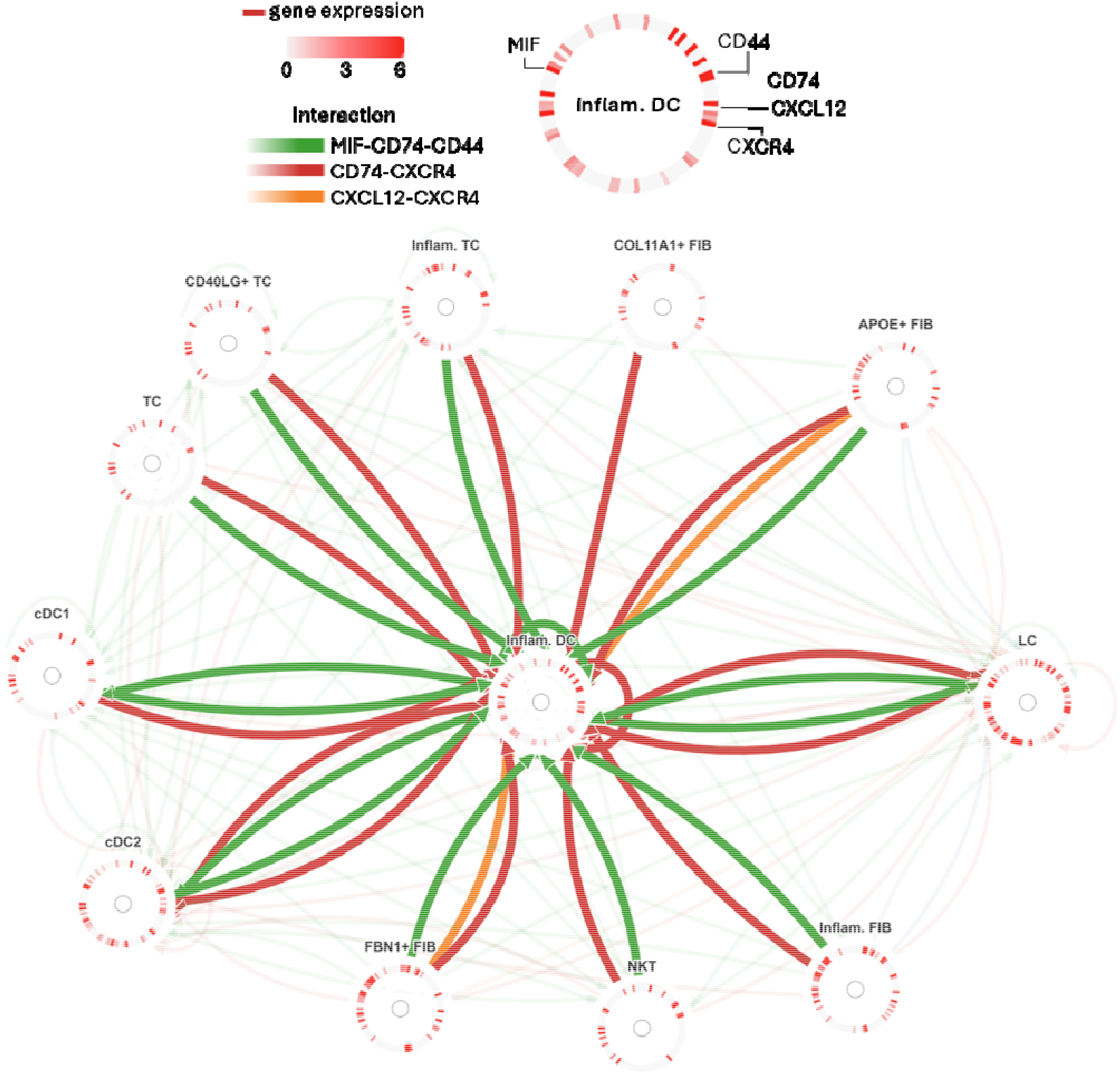
Cell communication network for atopic dermatitis. Each node represents a cell type and contains a ring. The ring’s sector elements represent the expression profiles of 106 genes across all cell types. Edges represent inferred cell-to-cell communications by CellChat. Edge color represents different ligand-receptor interactions for intercellular communication, while edge intensity represents the strength of communication between cells. The highlighted interactions are cellular communication between inflammatory DCs and other cell types with the highest scores. The ring annotation shows the location of the communication molecules (i.e., MIF, CD44, CD74, CXCR4, and CXCL12) in the ring. The genes’ descriptions can be found in Supplementary Table S2.

## Discussion and Conclusion

RingNet supports the flexible visualization of multimodal biological networks but lacks direct interoperability with Cytoscape’s extensive plugin ecosystem, such as network topology analysis, enrichment analysis, and systems biology workflows (Shannon *et al*. 2003). Specifically, RingNet does not yet support the bidirectional import or export of Cytoscape session formats (i.e., cys files). A major technical challenge is that RingNet uses a custom JSON structure optimized for compact, multi-ring visualization and aligned, multi-modal rendering. In contrast, Cytoscape relies on a different internal graph and attribute representation. Consequently, direct format conversion is nontrivial and would require a significant redesign of the serialization and rendering pipeline. Future development may explore partial interoperability through standard graph exchange formats to facilitate integration with Cytoscape plugins and downstream analytical workflows.

One limitation of RingNet is that its compact, ring-based representation can appear visually cluttered when displaying many data modalities simultaneously. While the framework supports multiple data layers, the addition of numerous independent encodings can increase visual complexity and reduce the interpretability of individual rings, especially in highly multimodal datasets. This is the reason we restrict the data type to five modalities rather than more in RingNet.

Another limitation of RingNet is its lack of native support for spatially resolved omics data, such as spatial transcriptomics and spatial multi-omics. Recent advances in spatial technologies have shown that maintaining tissue architecture and spatial neighborhood information is essential for understanding cellular heterogeneity, cell-cell communication, and disease microenvironments (Jain and Eadon 2024, Gulati *et al*. 2025). Spatial transcriptomics is increasingly integrated with transcriptomic, epigenomic, proteomic, and imaging modalities to characterize tissue organization and signaling networks in situ (Kiessling and Kuppe 2024, Liu X *et al*. 2024). However, in its current implementation, RingNet focuses on topology-based network visualization and does not incorporate spatial coordinates, tissue images, or spatial neighborhood constraints into the visualization framework. Future extensions could integrate spatial coordinates as node attributes or layered tissue maps to support the visualization of spatially informed interaction networks and multi-modal tissue architectures.

RingNet provides an interactive framework for visualizing multi-modal data within networks, enabling intuitive user-driven exploration. Its core strength lies in the compact, ring-based representation of multiple data layers at the sector element level (e.g., genes, samples, or cell groups), ensuring consistent alignment across modalities within each network node. This design allows users to directly assess correlations among diverse data types while simultaneously comparing distributional patterns across nodes or predefined groups. Such alignment supports clear interpretation of intra-sample relationships and reveals group-level heterogeneity, stratification effects, and coordinated shifts across data layers. Case studies in breast cancer genome-wide omics and atopic dermatitis single-cell communication networks demonstrate RingNet’s ability to uncover meaningful biological patterns and relationships. Beyond these examples, the framework can integrate computational analysis outputs, such as model-derived feature importance scores, alongside experimental measurements to support transparent interpretation and hypothesis generation. Overall, RingNet lowers barriers to exploring and communicating complex network-based biological insights.

## Availability and requirements

Project name: RingNet

Project home page: https://ringnet.rd.tuni.fi

Operating system(s): Web based tool

Programming language: R and JavaScript

Other requirements: Not required

License: GPL

Any restrictions to use by non-academics: No restrictions

**Ethics, Consent to Participate, and Consent to Publish declarations**: not applicable.

## Data availability

The RingNet is available at https://ringnet.rd.tuni.fi, which also hosts all datasets used in this study. The source code for RingNet is accessible at https://github.com/laixn/ringnet.

## Author contributions statement

**LZ**: methodology; software; data curation; writing – original draft; writing – review and editing. **XL**: conceptualization; methodology; software; data curation; investigation; validation; formal analysis; supervision; funding acquisition; visualization; project administration; resources; writing – original draft; writing – review and editing.

## AI usage declaration

After drafting the manuscript, we used ChatGTP 5.6 for language checking and grammar corrections. We are fully responsible for text refined by the AI-writing tool. We reviewed and edited the output as needed and take full responsibility for the content of the published article.

## Acknowledgments

We would like to express our gratitude to the anonymous reviewers who have offered valuable insights and constructive feedback on our paper since its initial publication as a preprint.

## Abbreviations

CNV: copy number variation
CSV: comma-separated value
HSTS: http strict transport security
HTML: hypertext markup language
HTTPS: hypertext transfer protocol secure
PNG: portable network graphics
SVG: scalable vector graphic
TCGA: the cancer genome atlas
TLS: transport layer security

