## Supplementary materials for "RingNet: An Interactive Platform for Multi-Modal Data Visualization in Networks"

### **Node-based pattern discovery**

We develop a method that identifies sample-level patterns in network nodes. This function helps users identify nodes with specific patterns in a network. In the case study presented in the main text Figure 3, we use synthetic data to demonstrate the function’ capability in identifying anti-correlated pattern in genes’ expression and DNA methylation profiles. Specifically, gene expression and methylation data are synthesized by estimating the distribution parameters of each gene's raw data. Gene expression is modeled in log space using a log-normal distribution because expression values are nonnegative and right-skewed. Methylation is modeled using a beta distribution because beta values are bounded between 0 and 1. The original tumor stage labels were used only as a sample-level progression index. Sample-level noise is added to mimic biological heterogeneity and patient-specific variation. In the following, we describe how the function is implemented in detail.

**Step1: Sample-level pattern hit identification**

The users first define thresholds for Data1 (D1) and Data2 (D2). For a gene (i), only the samples (s) satisfying the thresholds are used for follow-up analysis. The threshold can be set using data’s raw value or normalized value such as the z-score or min-max method. Once a sample is selected it is counted as supporting a pattern of interest, we then use its corresponding data’s z-score values to compute its score (Intensity). z-score keeps intensity measurement comparable across genes.

$${Hit}_{<g,i>}=\left\{ \begin{aligned} 1, if\left( {D1}_{<g,i>}\geq\tau_{<D1,H>}\cap{D2}_{<g,i>}\leq\tau_{<D2,L>} \right.) \\ 1, if \left( {D1}_{<g,i>}\leq\tau_{<D1,L>}\cap{D2}_{<g,i>}\geq\tau_{<D2,H>}) \right. \\ 0, if else \end{aligned} \right.$$

$${Intensity}_{<g,i>}=\left\{ \begin{aligned} \left| {Z_{D1}}_{<g,i>}\times{Z_{D2}}_{<g,i>} \right|, {if Hit}_{<g,i>}=1 \\ 0,{if Hit}_{<g,i>}=0 \end{aligned} \right.$$

τ_<{D1,D2},{H,L}>_ denotes the predefined high and low threshold for D1 and D2, respectively. Z_D1_ and Z_D2_ denote the z-score transformation for the data. Changing the threshold for each data can determine which samples support a pattern of interest. We are interested in identifying samples with anti-correlated patterns between D1 and D2 (i.e., negative correlation between gene expression and DNA methylation). Therefore, we select samples with greater value in D1 and smaller value in D2 or the opposite trend. Furthermore, the selected samples’ intensity score is computed using the data’s absolute z-score values and the other samples’ scores are set zero.

**Step 2: Group-level evidence aggregation**

When samples’ group annotation is available, we summarize sample-level hits within each group (j = 1…n). This group can represent cancer stage, ordered clinical group, time point or pseudotime bin. The group-specific HitRate measures how many samples in a group support the pattern of interest. IntensityScore averages z-score intensity among those supporting samples. These two quantities are combined into GroupIntensity so that a gene supported by many samples with moderate IntensityScore is distinguished from a gene supported by only a few samples with large IntensityScore values.

${HitRate}_{<g,j>} = \underset{i \in group j=1\ldots n}{\mathrm{mean}} \left( {Hit}_{<g,i>} \right.$)

$${IntensityScore}_{<g,j>} = \underset{i \in group j=1\ldots n}{\mathrm{mean}} ({Intensity}_{<g,i>})$$

${GroupIntensity}_{<g,j>} = \sqrt{{HitRate}_{j}\times{IntensityScore}_{j}},group j=1\ldots n$

GroupIntensity is a group-level signal. A larger value of GroupIntensity means that, within a group, the anti-correlated pattern between D1 and D2 is supported by a larger fraction of samples and/or has stronger z-score intensity. It indicates the negative correlation between the two data sets becomes stronger across the groups. Biologically, an increase in GroupIntensity from early to late tumor stages (characterized by group annotation) means that the anti-correlated pattern between gene expression and DNA methylation becomes more evident during disease progression.

**Step 3: TrendScore characterizes group-specific trend**

The first score, TrendScore, determines group specificity and can be used to interpret tumor progression when the groups are arranged in order of increasing tumor stages in the samples. This score contains two terms. The first, TrendIntensity, quantifies anti-pattern strength along the ordered groups. The second, AntiTrend, quantifies the correlation between the average D1 and D2 profiles across the groups. We calculate both terms using Spearman's rank correlation to quantify the correlation coefficient between groups (e.g., tumor stage) and their corresponding molecular profiles. These profiles include GroupIntensity, D1 (i.e., the average gene expression profile), and D2 (i.e., the average DNA methylation profile). A higher value indicates that the intensity increases and the two molecular profiles move in opposite directions from early to late tumor stages.

$${Trend}_{<g,Inensity>} = \mathrm{corr} \left( Group[1\ldots n], \left[ {GroupIntensity}_{<g,1>}\ldots{GroupIntensity}_{<g,n>} \right]) \right.$$

$${Trend}_{<g,D1>}= \mathrm{corr} \left( Group[1\ldots n], \left[ {mean(D1)}_{<g,1>}\ldots{mean(D1)}_{<g,n>} \right]) \right.$$

$${Trend}_{<g,D2>}= \mathrm{corr} \left( Group[1\ldots n], \left[ {mean(D2)}_{<g,1>}\ldots{mean(D2)}_{<g,n>} \right]) \right.$$

$${AntiTrend}_{g}=\max\left( 0, {Trend}_{<g,D1>}) \right.\times\max\left( 0, -{Trend}_{<g,D2>} \right.) + \max\left( 0, -{Trend}_{<g,D1>}) \right.\times\max\left( 0, {Trend}_{<g,D2>}) \right.$$

$${TrendScore}_{g} = 0.5\times max \left( 0, {Trend}_{<g,Inensity>}) \right. + 0.5 {\times AntiTrend}_{g}$$

In our case, a gene whose GroupIntensity increases with tumor stage is given more weight than a gene with similar values across all stages. Furthermore, AntiTrend requires that the stage-level averages of the two molecular layers be directionally opposite. Thus, TrendScore prioritizes genes whose anti-pattern is detectable and organized in a manner resembling tumor progression.

**Step 4: GroupScore determines group-level intensity supporting the trend**

The second score, GroupScore, averages GroupIntensity across all groups and quantifies the amount of group-level evidence supporting the pattern of interest. Importantly, this score is not weighted along tumor-stage progression because the tendency specific to each group is already captured by TrendIntensity.

$${GroupScore}_{g} = \underset{i \in group j}{\mathrm{mean}} \left( {GroupIntensity}_{<g,i>}) \right.$$

The score represents the average amount of reliable anti-correlated pattern signals across stages. A higher GroupScore indicates that more samples support z-score-based anti-pattern intensity across the cohort. A low GroupScore indicates that the trend may be based on fewer samples. Thus, GroupScore prevents weak but visually monotonic patterns from being overinterpreted.

**Step 5: Node filtering using one or two thresholds**

When using both TrendScore and GroupScore, a node (i.e., a gene in a network) is retained if it passes both user-defined thresholds.

$${Retain}_{g}=\left\{ \begin{aligned} Yes, if\left( {TrendScore}_{g} \geq\tau_{T}\cap{GroupScore}_{g} \geq\tau_{G} \right.) \\ No, if else \end{aligned} \right.$$

Specifically, τ_T_ and τ_G_ denotes the TrendScore and GroupScore thresholds, respectively. Both thresholds can be predefined by users at the frontend. Increasing τ_T_ makes pattern discovery more selective for nodes whose GroupIntensity profile has a stronger group-associated trend. Increasing τ_G_ makes pattern discovery more selective for nodes whose pattern evidence is supported by stronger and/or more consistent stage-level evidence. These two thresholds therefore control two different dimensions of biological stringency: τ_T_ controls tumor progression specificity, whereas τ_G_ controls sample sufficiency.

In addition, if only GroupScore was used, RingNet would preferentially retain genes with strong anti-pattern evidence, even when that evidence is present uniformly across all groups and is not progression specific. Such genes may be biologically meaningful as anti-pattern nodes across groups. Conversely, if only TrendScore was used, genes with an attractive monotonic trend but weak sample-level support could be retained. Requiring both thresholds ensures that the pattern discovery function prioritizes nodes with both a group-associated trend and sufficient samples supporting the observed trend.

**Heterogenous and homogeneous sample identification in a node**

We use a method that characterizes the distance between samples by considering the directional and magnitude differences in their profiles. This function helps users identify samples with heterogeneous or homogeneous profiles in a network node.

**Sample vector representation**

Two four-dimensional vectors are constructed for each sample s. The global vector G_s_ is the average value of a dataset across all samples. The local vector L_s_ is the actual value of the current sample. Each sample has four-omics data including gene expression (exp), DNA methylation (mty), copy number variation (cnv), and single nucleic tide variation (snv).

$$G_{s} = [G_{exp} \left( s \right), G_{mty} \left( s \right), G_{cnv} \left( s \right), G_{snv} \left( s \right)]$$

$$L_{s} = [L_{exp} \left( s \right), L_{mty} \left( s \right), L_{cnv} \left( s \right), L_{snv} \left( s \right)]$$

$$Z_{s} = 0.60 \times G_{s}+0.40\times L_{s}=$$

$$[0.60\times G_{exp} \left( s \right) + 0.40{\times L}_{exp} \left( s \right), 0.60 \times G_{mty} \left( s \right)+ 0.40 \times L_{mty} \left( s \right),$$

$$0.60 \times G_{cnv} \left( s \right) + 0.40\times L_{cnv} \left( s \right), 0.60 \times G_{snv} \left( s \right) + 0.40 \times L_{snv} \left( s \right)]$$

Z_s_ is a four-dimensional integrated vector. The 0.60 weight emphasizes network-level evidence, and the 0.40 weight preserves node-specific local evidence.

**Sample distance calculation**

For any two samples s_i_ and s_j_, we use the mixed distance (D_mix_) that is computed from samples’ integrated vectors in the following steps. First, cosine similarity is computed between the integrated vectors:

$\mathrm{CosSim} \left( s_{i}, s_{j} \right) = \frac{Z_{s_{i}} \cdot Z_{s_{j}}}{\left\| Z_{s_{i}} \right\|\times\left\| Z_{s_{j}} \right\|}$ with

$Z_{s_{i}} \cdot Z_{s_{j}}=\sum_{k=1}^{n} Z_{s_{i}} \cdot Z_{s_{j}},\left\| Z_{s_{i}} \right\|=\sqrt{{\sum_{k=1}^{n} Z_{s_{i}}}^{2}}$ , and $\left\| Z_{s_{j}} \right\|=\sqrt{{\sum_{k=1}^{n} Z_{s_{j}}}^{2}}$

Where n is the total number of features used for computing the distance. In our case, n = 4 x g, where g is the number of genes in each sample and each gene has four types of data. Cosine similarity is then converted to a cosine distance in the range [0,1]:

$$D_{cos} \left( s_{i}, s_{j} \right) =\frac{1 -\mathrm{CosSim}\left( s_{i}, s_{j} \right)}{2}$$

Second, the normalized Euclidean distance scales the L2 distance by the maximum pairwise distance observed within the current network (N):

$$D_{euc} \left( s_{i}, s_{j} \right) =\frac{\left\| Z_{s_{i}} - Z_{s_{j}} \right\|^{2}}{\max_{(p, q) \in N} \left\| Z_{p} - Z_{q} \right\|^{2}}$$

p and q are samples having the maximum Euclidean distance within the network.

Finally, the two distance components are combined into the mixed distance D_mix_:

$$D_{mix} \left( s_{i}, s_{j} \right) = 0.60\times D_{euc} \left( s_{i}, s_{j} \right) + 0.40{\times D}_{cos} \left( s_{i}, s_{j} \right)$$

D_cos_ captures directional difference and D_euc_ captures magnitude difference. So, D_mix_ reflects both signal intensity separation and multi-omics pattern direction. D_mix_ is used to identify heterogenous and homogeneous samples in a network node.

**Methods for identifying different and similar samples**

When group annotations are provided (group-aware mode), samples are first separated by groups. For each group k, the group-specific sample pool S_k_ and the capped display count C_k_ are defined as follows:

$$S_{k}=\left\{ s_{i} \mid\mathrm{group} \left( s_{i} \right) = k \right\}$$

$$C_{k}=\min\left( m, S_{k} \right)$$

Where m is a number that can be adjusted by users, and we set it to 8.

**Difference displaying reveals heterogeneous and divergent samples.** For each group k, difference displaying selects up to C_k_ samples that are mutually far apart based on the computed D_mix_. The selected difference set is A_k_. It is initialized using the most distant pair:

$$\left( a_{1}, a_{2} \right)=\underset{s_{i}, s_{j} \in S_{k}}{arg max}D_{mix} \left( s_{i}, s_{j} \right),\quad A_{k}=\{a_{1}, a_{2}\}$$

Then each new sample is selected by the max-min rule. This chooses the candidate whose nearest selected neighbor is still as far away as possible, so the displayed samples spread across separated multi-omics states:

$$s^{*}=\underset{s \in S_{k}\backslash A_{k}}{arg max}\min_{u \in A_{k}}D_{mix} \left( s, u \right)$$

**Similarity displaying reveals homogeneous sample groups.** For each group k, similarity displaying selects up to C_k_ samples that are mutually close according to D_mix_. The selected similarity set is B_k_. It is initialized using the closest pair:

$$\left( b_{1}, b_{2} \right) = \underset{s_{i}, s_{j} \in S_{k}}{arg min}D_{mix} \left( s_{i}, s_{j} \right),\quad B_{k} = \{b_{1}, b_{2}\}$$

Then each new sample is selected by minimizing its average distance to the already selected group.

$$s^{*}=\underset{s \in S_{k}\backslash B_{k}}{arg min} \underset{u \in B_{k}}{\mathrm{mean}} D_{mix} \left( s, u \right)$$

If no group information is available (group-free mode), all samples are treated as one sample pool (S_all_ = {S_1_,…,S_n_} and C = min(m, S_all_)). Then, the same methods can be applied to identify heterogeneous and homogeneous samples.

In both group-aware and group-free modes, selection outputs global sample indices, not node-specific local positions. The same index must refer to the same sample in four data sets for every node. Thus, selection changes which samples are displayed, but it does not reorder where samples sit in the node ring. Both modes preserve alignment across all four omics rings and across all nodes.

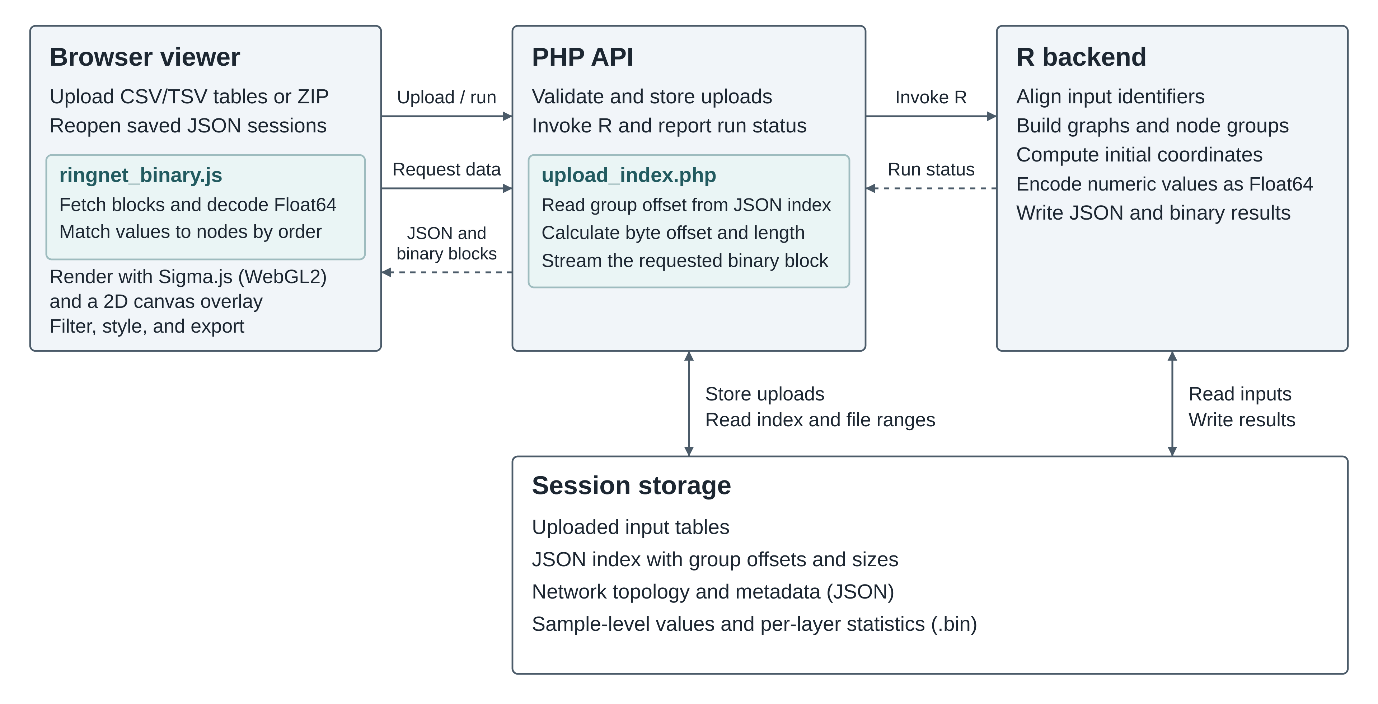

**Figure S1. RingNet system architecture and data flow.** The browser viewer accepts numbered input tables or a ZIP archive and can reopen a saved session JSON. The PHP API validates and stores uploads, invokes R, and serves indexed results. R aligns identifiers, builds the graph and its node-group views, and computes initial coordinates. Session storage holds the uploaded inputs, indexed JSON for network topology and metadata, and binary blocks for sample-level numeric values.

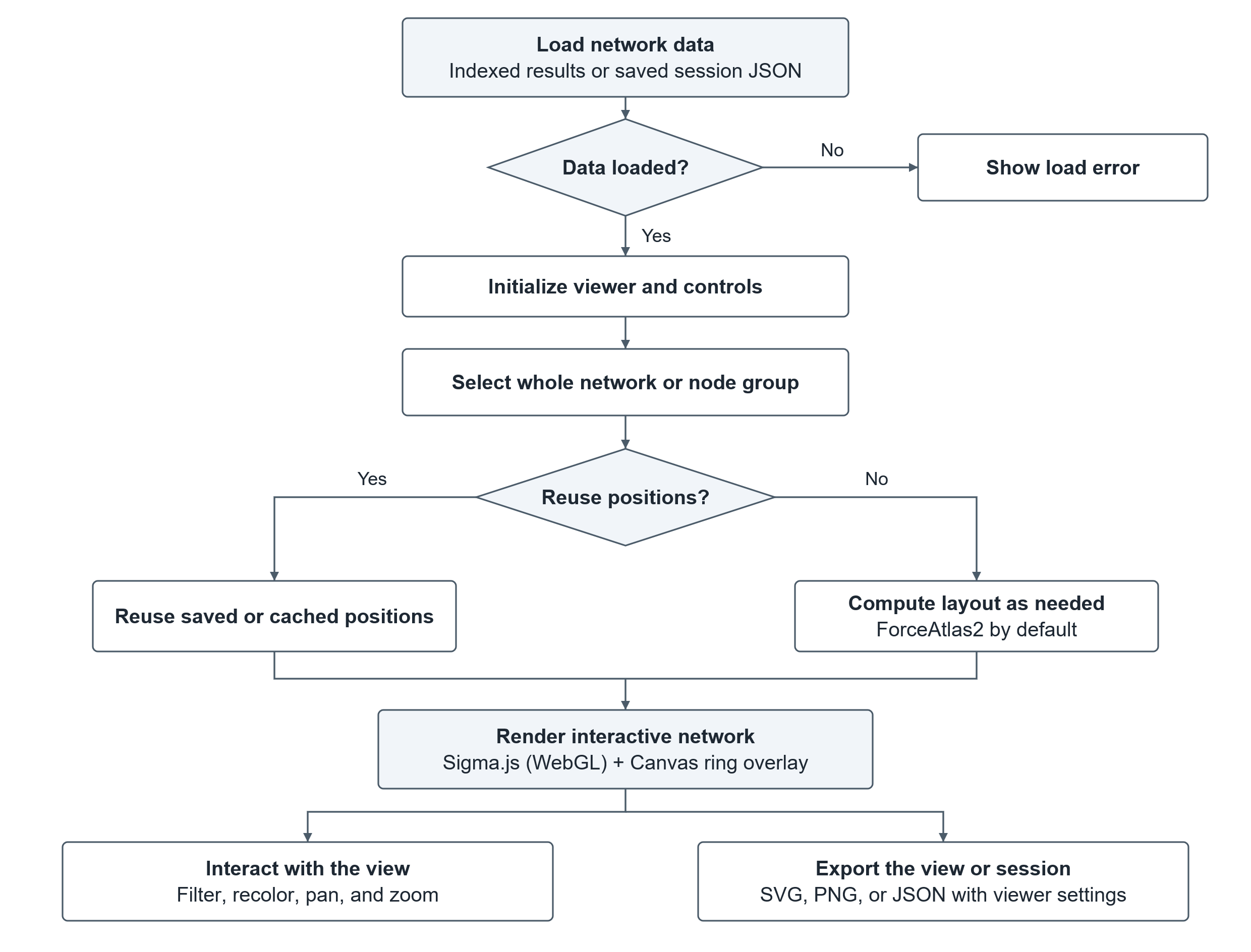

**Figure S2. RingNet frontend workflow.** The viewer loads indexed network data or opens a saved session JSON and initializes its controls. After a network is selected, the viewer reuses stored positions when available or computes a layout. New networks use ForceAtlas2 in a background worker by default. Sigma.js renders the graph with WebGL, and a Canvas overlay draws the rings. Users can filter, recolor, pan, and zoom, export SVG or PNG images, and save a session JSON containing viewer settings.

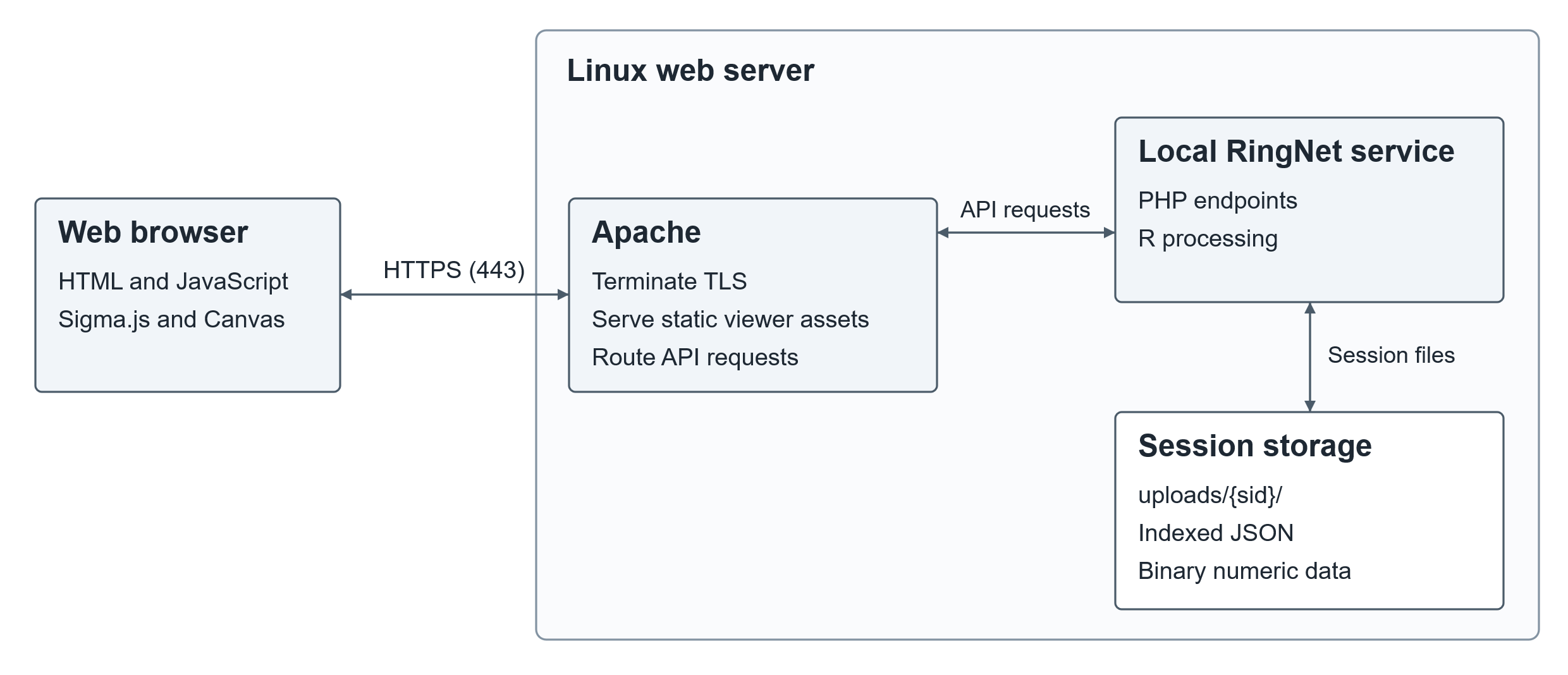

**Figure S3. RingNet deployment and request routing.** The browser connects over HTTPS to Apache on the Linux web server. Apache terminates TLS, serves viewer assets, and routes API requests to the local RingNet service. The PHP endpoints invoke R and deliver indexed JSON and binary numeric data from the session directory. HSTS is configurable on the web server.

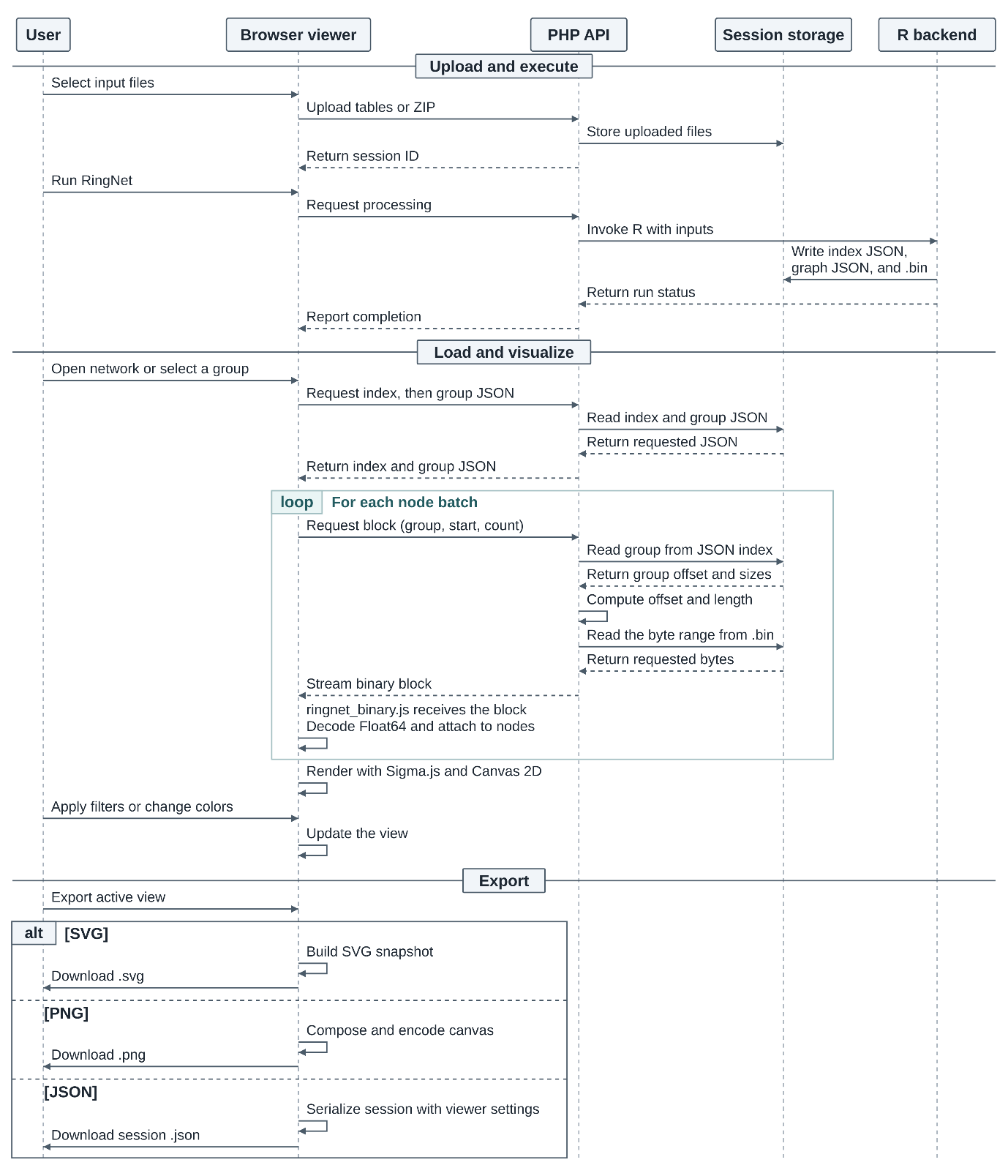

**Figure S4. RingNet’s end-to-end workflow.** The diagram traces input upload, R execution, result retrieval, interactive visualization, and export. The browser uses a session identifier to request network data through the PHP API. Users can export the active network view as SVG or PNG or download a session JSON containing viewer settings.

**Table S1** the table shows the computational resources used to visualize 25 networks simultaneously. The columns are the network indexes, the number of nodes in each network, the number of edges in each network, the computation time (in seconds) used by RingNet, and the amount of RAM (in megabytes) used by RingNet, including the resident set size (RSS) and the high-water mark (HWM). rss_start_mb: The amount of RAM the process was using at the beginning. rss_end_mb: The amount of RAM used at the end. rss_diff_mb: The net difference in RSS in RAM usage. hwm_start_mb: The peak reached before the tracking period began. hwm_end_mb: The new peak reached by the end of the process. hwm_diff_mb: The difference in the HWM amount by which the previous peak was exceeded. The identical rss_end_mb and hwm_end_mb suggest that the process was at its maximum memory consumption right when it finished. There was no "spike" in the middle that was higher than the final value; the memory usage likely climbed steadily or jumped and stayed at that new level.

| net_index | net_node | net_edge | time_sec | rss_start_mb | rss_end_mb | rss_diff_mb | hwm_start_mb | hwm_end_mb | hwm_diff_mb |
| --- | --- | --- | --- | --- | --- | --- | --- | --- | --- |
| 1 | 169 | 247 | 2.06 | 202.93 | 268.08 | 65.16 | 202.93 | 268.08 | 65.16 |
| 2 | 1821 | 5106 | 21.72 | 268.08 | 395.07 | 126.99 | 268.08 | 395.07 | 126.99 |
| 3 | 635 | 2235 | 3.61 | 395.07 | 397.39 | 2.32 | 395.07 | 397.39 | 2.32 |
| 4 | 167 | 306 | 1.62 | 397.39 | 397.39 | 0.00 | 397.39 | 397.39 | 0.00 |
| 5 | 1875 | 6100 | 15.95 | 203.22 | 381.48 | 178.26 | 203.22 | 381.48 | 178.26 |
| 6 | 138 | 167 | 1.58 | 381.48 | 381.48 | 0.00 | 381.48 | 381.48 | 0.00 |
| 7 | 111 | 150 | 1.55 | 381.48 | 381.48 | 0.00 | 381.48 | 381.48 | 0.00 |
| 8 | 109 | 185 | 1.90 | 203.28 | 268.59 | 65.31 | 203.28 | 268.59 | 65.31 |
| 9 | 445 | 1099 | 3.43 | 268.59 | 313.38 | 44.80 | 268.59 | 313.38 | 44.80 |
| 10 | 91 | 96 | 1.45 | 313.38 | 313.38 | 0.00 | 313.38 | 313.38 | 0.00 |
| 11 | 90 | 79 | 1.41 | 313.38 | 313.38 | 0.00 | 313.38 | 313.38 | 0.00 |
| 12 | 83 | 60 | 1.78 | 203.14 | 268.23 | 65.09 | 203.14 | 268.30 | 65.17 |
| 13 | 95 | 116 | 1.65 | 268.23 | 301.24 | 33.02 | 268.30 | 301.24 | 32.94 |
| 14 | 84 | 61 | 1.73 | 301.24 | 305.63 | 4.38 | 301.24 | 305.63 | 4.38 |
| 15 | 126 | 187 | 1.90 | 203.29 | 268.59 | 65.31 | 203.29 | 268.59 | 65.31 |
| 16 | 87 | 772 | 1.57 | 268.59 | 300.83 | 32.24 | 268.59 | 300.83 | 32.24 |
| 17 | 72 | 38 | 1.51 | 300.83 | 304.18 | 3.35 | 300.83 | 304.18 | 3.35 |
| 18 | 87 | 59 | 1.51 | 304.18 | 307.54 | 3.35 | 304.18 | 307.54 | 3.35 |
| 19 | 69 | 405 | 1.80 | 203.12 | 268.27 | 65.16 | 203.12 | 268.30 | 65.19 |
| 20 | 77 | 46 | 1.56 | 268.27 | 299.79 | 31.52 | 268.30 | 299.79 | 31.48 |
| 21 | 62 | 30 | 1.49 | 299.79 | 299.79 | 0.00 | 299.79 | 299.79 | 0.00 |
| 22 | 52 | 20 | 1.72 | 203.27 | 268.29 | 65.02 | 203.27 | 268.37 | 65.10 |
| 23 | 61 | 27 | 1.43 | 268.29 | 268.81 | 0.52 | 268.37 | 268.81 | 0.44 |
| 24 | 123 | 119 | 1.65 | 268.81 | 301.87 | 33.06 | 268.81 | 301.87 | 33.06 |
| 25 | 98 | 70 | 1.54 | 301.87 | 307.28 | 5.41 | 301.87 | 307.28 | 5.41 |

**Table S2** The table lists the full names of the genes visualized in the networks in Figure 4, 5, and 6 in the main text.

| Gene symbol – Figure 4 | Full name |
| --- | --- |
| ADRB1 | adrenoceptor beta 1 |
| C5AR1 | complement C5a receptor 1 |
| GNA14 | G protein subunit alpha 14 |
| GNAI2 | G protein subunit alpha i2 |
| HRH1 | histamine receptor H1 |
| HTR4 | 5-hydroxytryptamine receptor 4 |
| KCNAB1 | potassium voltage-gated channel subfamily A regulatory beta subunit 1 |
| NUMR1 | neuromedin U receptor 1 |
| Gene symbol – Figure 5 | **Full name** |
| ADRA2A | Adrenoceptor alpha 2A |
| ADRB2 | Adrenoceptor beta 2 |
| AGTR1 | Angiotensin II receptor type 1 |
| BDKRB2 | Bradykinin receptor B2 |
| GNA11 | G protein subunit alpha 11 |
| GNA12 | G protein subunit alpha 12 |
| GNA14 | G protein subunit alpha 14 |
| GNAI1 | G protein subunit alpha i1 |
| GNAQ | G protein subunit alpha q |
| GNAZ | G protein subunit alpha z |
| GNB5 | G protein subunit beta 5 |
| GNG2 | G protein subunit gamma 2 |
| GRK5 | G protein-coupled receptor kinase 5 |
| HRH2 | Histamine receptor H2 |
| PTGER3 | Prostaglandin E receptor 3 |
| S1PR1 | Sphingosine-1-phosphate receptor 1 |
| TACR1 | Tachykinin receptor 1 |
| Gene symbol – Figure 6 | **Full name** |
| CD44 | CD44 molecule (Indian blood group) |
| CD74 | CD74 molecule |
| CXCL12 | C-X-C motif chemokine ligand 12 |
| CXCR4 | C-X-C motif chemokine receptor 4 |
| MIF | Macrophage migration inhibitory factor |
